# DNA damage response in peripheral mouse blood leukocytes *in vivo* after variable, low-dose rate exposure

**DOI:** 10.1101/648568

**Authors:** Qi Wang, Monica C. Pujol, Guy Garty, Maria Taveras, Jay Perrier, Carlos Bueno-Beti, Igor Shuryak, David J. Brenner, Helen C. Turner

**Author notes:** Corresponding author: Qi Wang.

## Abstract

Environmental contamination and ingestion of the radionuclide Cesium-137 (^137^Cs) is a large concern in fallout from a nuclear reactor accident and improvised nuclear device and highlights the need to develop biological assays for low dose rate, internal emitter radiation. To mimic low dose rates attributable to fallout, we have developed a VAriable Dose-rate External ^137^Cs irradiatoR (VADER), which can provide arbitrarily varying and progressive low dose rate irradiations in the range of 1.2 to 0.1 Gy/day, while circumventing the complexities of dealing with radioactively-contaminated biomaterials. We investigated the kinetics of mouse peripheral leukocytes DNA damage response in vivo after variable, low-dose rate ^137^Cs exposure. C57BL/6 mice were placed in the VADER over 7 days with total accumulated dose up to 2.7 Gy. Peripheral blood response including the leukocytes depletion, apoptosis signal protein p53 and DNA repair biomarker γ-H2AX were measured. The results illustrated that blood leukocyte count had significantly dropped by days 7. P53 levels peaked at day 2 (total dose=0.91Gy) and then declined whereas γ-H2AX yields generally increased with accumulated dose and peaked at day 5 (total dose=2.08Gy). ROC curve analysis for γ-H2AX provided a good discrimination of accumulated dose < 2 Gy and ≥ 2 Gy, highlighting the potential of γ-H2AX as a biomarker dosimetry in a protracted, environmental exposure scenario.

## Introduction

The effect of internal emitters and associated varying dose rate effects on biodosimetry are not well studied. Cesium-137 (^137^Cs), a fission product of uranium and plutonium is a radionuclide of concern and major environmental contaminant following the detonation of an improvised nuclear device (IND) or a nuclear reactor accident (Garty et al. 2017; Simon et al. 2004; Yasunari et al. 2011). In this exposure scenario, a significant amount of soluble radionuclides may be dispersed into the atmosphere as a component of fallout, which could affect vast numbers of people located at substantial distances from the radiological event site. The Chernobyl and Fukushima Daiichi accidents both illustrate the large-scale detrimental effects of the environmental radioactive contamination (López-Vicente et al. 2018; Moysich et al. 2002; Nakamura et al. 2017; Steinhausler 2005). Thus, there is a need to develop high-throughput biodosimetry systems and validate biomarkers for protracted ionizing radiation in population-based triage following a large-scale accidental or malicious release of radioactive substances.

Compared with external exposure, the dose rates of internal exposures are typically much lower (below 1 Gy/day). At the Columbia University Center for High Throughput Minimally Invasive Radiation Biodosimetry in Center Medical Countermeasures against Radiation (CMCR), we have used a ^137^CsCl internal emitter mouse model to examine the long-term effects of ^137^CsCl activity for cytogenetic (Turner et al. 2015), transcriptomic (Paul et al. 2014) and metabolomic (Goudarzi et al. 2013) endpoints. The challenge with using the mouse ^137^Cs injection studies is that all the biomaterials are radioactively contaminated, and as such, require dedicated equipment, costly clean-up and decontamination procedures. To overcome this drawback, we have developed a VAriable Dose-rate External ^137^Cs irradiatoR (VADER) to facilitate modeling of low dose rate ^137^Cs exposure in mice using external irradiations. Based on the repurposing of using retired ^137^Cs brachytherapy seeds, the VADER irradiator is designed to provide arbitrarily varying and constant low dose rate irradiations in the range 1.2 to 0.1 Gy/day, resulting in chronic exposures of Gy-level doses over days to weeks of exposure (Garty et al. 2019a; Garty et al. 2017). The advantages of such a system are that it allows for experiments in a controlled, non-radioactive setting which can be programmed to mimic internal ^137^Cs biokinetics.

In our earlier work, we showed the persistence of radiation-induced DNA γ-H2AX double strand breaks (DSBs) *in vivo*, several weeks after the administration of ^137^Cs internal emitter gamma radiation which is in clear contrast to acute dose exposures where the γ-H2AX is typically decayed after 24-48 h due to DSB repair (Halm et al. 2014; Turner et al. 2019; Turner et al. 2010; Turner et al. 2015). We injected C57Bl/6 mice with a range of ^137^CsCl activities to achieve total-body committed doses of ~ 4 Gy at days 3, 5, 7, and 14 of ^137^Cs exposure. Despite the complicated nature of the studied biological system and the time-dependent changes in radiation dose and dose rate due to radionuclide excretion, we were able to profile γ-H2AX repair kinetics and develop a semi-empirical model for predicting initial ^137^Cs incorporation activity, based on measurements 2-5 days after initial incorporation (Turner et al. 2019).

The objective of the present work was to extend our mouse ^137^Cs injection studies and measure DNA damage response in peripheral blood leukocytes using lower dose/dose rates which was not previously tested. The study was designed to expose C57BL/6 mice to VADER irradiation over 7 days (dose rate < 0.5 Gy/day) and accumulated doses ranging from 0.49 Gy (day 1) to 2.68 Gy (day 7). We measured the following endpoints: 1) mouse leukocytes depletion, 2) apoptosis levels, 3) p53 expression and 4) γ-H2AX kinetics. We present here a γ-H2AX assay based on imaging flow cytometry (Lee et al. 2019) to measure DNA damage in mouse blood leukocytes *in vivo* after progressively decreasing low dose rate external exposures.

## Materials and Methods

### Experimental animals

The animal studies were approved by the approved by the Columbia University Institutional Animal Care and Use Committee (IACUC, #AC-AAAQ2410). Male C57BL/6 mice (approximately 7 weeks old, 20-30 g) were purchased from Charles River Laboratories (Frederick, MD) and quarantined for a minimum of 14 days prior to group assignment by body weight stratification for randomization onto the study.

### Irradiations and dosimetry

#### i) The variable dose-rate external ^137^Cs irradiator (VADER)

The VADER is a custom built irradiation system, built to deliver controlled dose rates in the range 0.1-1 Gy/day to a cohort of up to 15 mice. The VADER uses ~0.5 Ci of retired ^137^Cs brachytherapy seeds, arranged in two platters, and placed above and below a “mouse hotel”. The sources can be moved under computer control to provide the required dose rate as a function of time. The VADER is shielded such that dose rate in the room is <0.1 mGy/wk.

#### ii) VADER mouse hotel setup

The hotel consists of a 35 cm × 35 cm × 12 cm acrylic box with sufficient bedding material, in which up to 15 mice are free to move and interact with each other. Within the hotel, mice are free to move around, eat and drink *ad libitum*. Temperature, humidity, air flow and lighting are fully controlled to the required animal care standards. All these environmental controls and monitoring are integrated into removable, mouse hotel so that they can be easily replaced in case of radiation damage. To monitor the environment in the mouse hotel, a temperature/humidity sensor (HWg HTemp, TruePath Technologies Victor, NY) is integrated into one of the hotel walls. Real time monitoring of the mice is performed using a 180° fisheye USB camera (ELP, Amazon) embedded into one of the hotel walls.

#### iii) Mouse TLD dosimetry

To ensure the dose that each mice received*, in vivo* dosimetry was performed on a mouse-by-mouse basis, by injecting a glass encapsulated TLD chip into each mouse (Garty et al. 2019a). Anesthesia was induced with 2% isoflurane delivered in 100% oxygen for <3 min before the implantation procedure. The encapsulated TLD rods (one per mouse) were placed in a 12-gauge needle coupled with a needle injector (Allflex, Irving, TX) and administered by subcutaneous injection in the dorsal neck. Following implantation mice were monitored up to 48 hours for complications. TLD rods were later read using a Harshaw 2500 TLD reader (Thermo Fisher Scientific™, Waltham, MA), most experiments used a heating profile consisting of a 5 °C/sec ramp up to 300 °C followed by a short hold at 300 °C and cool down to 50 °C. Dose was reconstructed based on the integrated light yield at a temperature higher than 180 °C to eliminate the low temperature, time dependent, glow peak (Garty et al. 2019a).

#### iv) VADER irradiation and dosimetry

The VADER was programmed to provide the dose rate profile for 2 batches of experiments shown in Fig. 1 with the source position updated hourly. This profile corresponds to the double exponential decay measured by Paul et al (Paul et al. 2014)

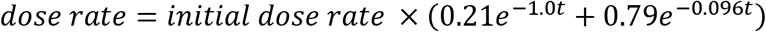

**Fig. 1.**
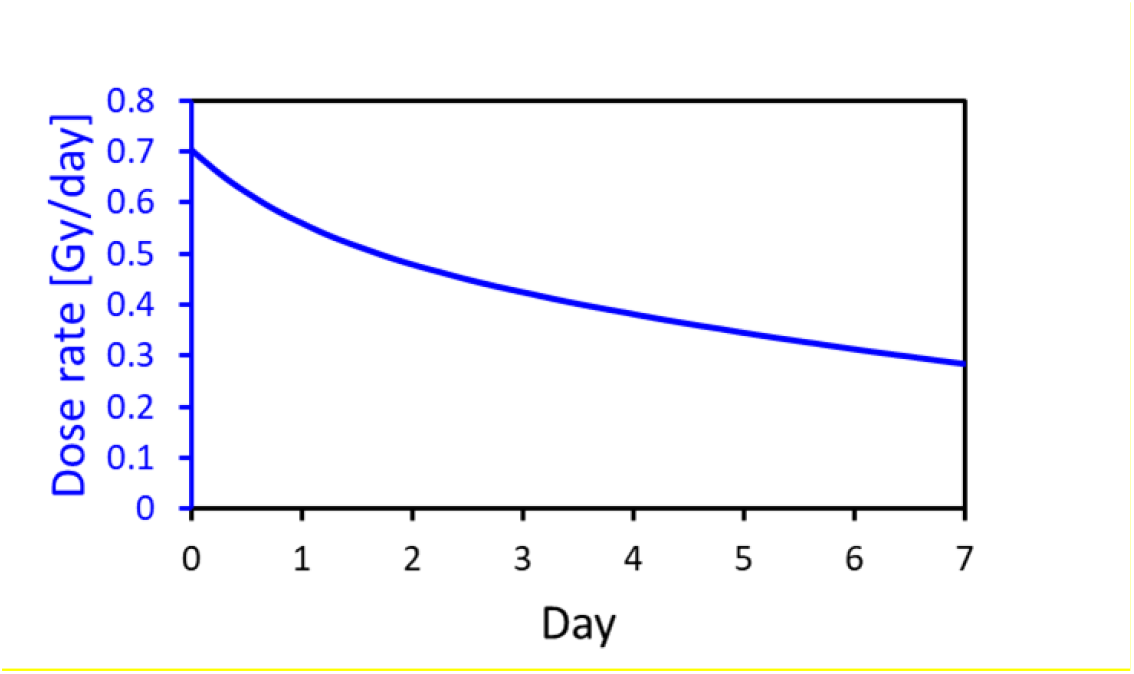
Dose rate vs time programmed into the VADER.

The dose rate was converted to source retraction using the VADER calibration curve:

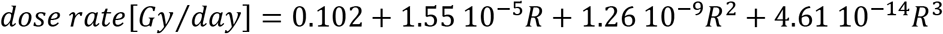

Where R is the source position, with *R=0* corresponding to the lowest dose rate of 0.1 Gy/day and *R=34,200 steps* corresponds to 1 Gy/day.

#### v) Mouse Irradiation

Two randomized batches of 15 mice were then loaded in the VADER hotel for irradiation respectively as the maximum number of mice that VADER can house is 15 mice. The first batch of mice were irradiated for 3 days, 4 days and 7 days. The second batch of mice were irradiated for 1 day, 2 days and 5 days. Randomized control mice were housed in an identical hotel with monitor system. For each time/dose point 5 irradiated mice and 4-5 control mice were assigned. Both the mice in the VADER hotel and control mice were monitored at least 4 times a day. The temperature in the VADER and control cage was 20°C to 25°C. Humidity in both ranged from 40% to 60%.

### Blood sample collection and cell counts

VADER-irradiated mice were sacrificed with paired non-irradiated sham-control mice on the same day after irradiation. All mice were euthanized by CO_2_ asphyxiation prior to blood collection. Peripheral whole blood samples were collected from each mouse by cardiac puncture using a heparin-coated syringe. Leukocyte, T cell and B cell counts were determined by flow cytometry using 20 μL of heparinized blood following the standard flow cytometry surface staining protocol (Wang et al. 2016). Peripheral blood was surface stained with APC/Cy7 conjugated anti-mouse CD45 antibody (clone 30-F11, Biolegend, San Diego, CA), APC conjugated anti-mouse CD3 antibody (clone 145-2C11, Biolegend) and PE conjugated anti-mouse CD19 antibody (clone 6D5, Biolegend), lysed with FACS lysing solution (BD Biosciences, Franklin Lakes, NJ) washed in phosphate-buffered saline (PBS, Gibco, Waltham, MA) with 5% fetal bovine serum (FBS) and measured on flow cytometry with well-established compensated matrix (CytoFLEX, Beckman Coulter, Pasedena, CA). Analyses were performed using CytExpert Software (Beckman Coulter).

### Imaging flow cytometry p53 and γ-H2AX analysis

Peripheral blood samples were lysed with RBC lysis buffer (Invitrogen^TM^ Waltham, MA), fixed using the FIX & PERM™ Cell Permeabilization Kit (Thermo Fisher Scientific™, Waltham, MA), washed with perm/wash buffer from the kit, and stained intracellularly by a rabbit polyclonal γ-H2AX (Abcam, Cambridge, MA) or a rabbit polyclonal p53 (Phospho-Ser37, Aviva systems biology, San Diego, CA). Proper isotype controls were included for intracellular staining as negative controls (Abcam, Rabbit polyclonal IgG). Following washing, cells were stained with a goat anti-rabbit Alexa Fluor 488 secondary antibody (Life technology, Carlsbad, CA). Samples were then washed with PBS, stained with nuclear dye DRAQ5 (Thermo Fisher Scientific™) and measured using the ImageStream^®X^ MkII Imaging Flow Cytometer (LUMINEX Corporation, Austin, Texas). Images of more than 5000 cells per sample were acquired at 40X magnification using the 488 nm excitation laser. For the compensation, cells stained with γ-H2AX antibody, p53 antibody or DRAQ5 only were captured using the 488 nm laser without with inactivated bright field illumination. The compensation coefficients were acquired automatically by the IDEAS 6.2 compensation wizard. Captured images were analyzed using IDEAS^®^ (LUMINEX Corporation, Austin, Texas) software for measuring proportions of apoptotic cell, γ-H2AX and p53 fluorescence intensity and γ-H2AX foci formation.

Fig. 2 shows the representative gating strategy to identify focused non-apoptotic cell population for measurement of fluorescence intensity of γ-H2AX and p53, and quantification of the γ-H2AX foci. The focused cells were gated according to the gradient similarity feature by visual inspection of cell images in the bright field channel (Fig. 2a). Single cells were then selected from images according to their area and aspect ratio in the bright field channel (Fig. 2b) and nucleated cells are selected based on DRAQ5 positivity to exclude the dead cells (Fig. 2c). As apoptotic cells showed a different pattern of γ-H2AX staining (Solier and Pommier 2014), they were then excluded according to the complexity of the nucleus and contrast of brightfield morphology (Fig. 2d). The mean fluorescence intensity (MFI) of γ-H2AX and p53 within the gated cells population was then analyzed and exported from the IDEAS^®^ software. The γ-H2AX foci formation was identified using the spot counting wizard (Fig. 2e). This wizard automatically creates masks based on peak mask identified intensity areas from an image with local maxima (bright) or minima (dark), followed by enumerate foci identified by the mask. Fig. 2f is a representative histogram for quantify foci number of each cell in one sample. A template file was generated and applied to all data files and automatically batch processed on IDEAS^®^.

**Fig. 2.**
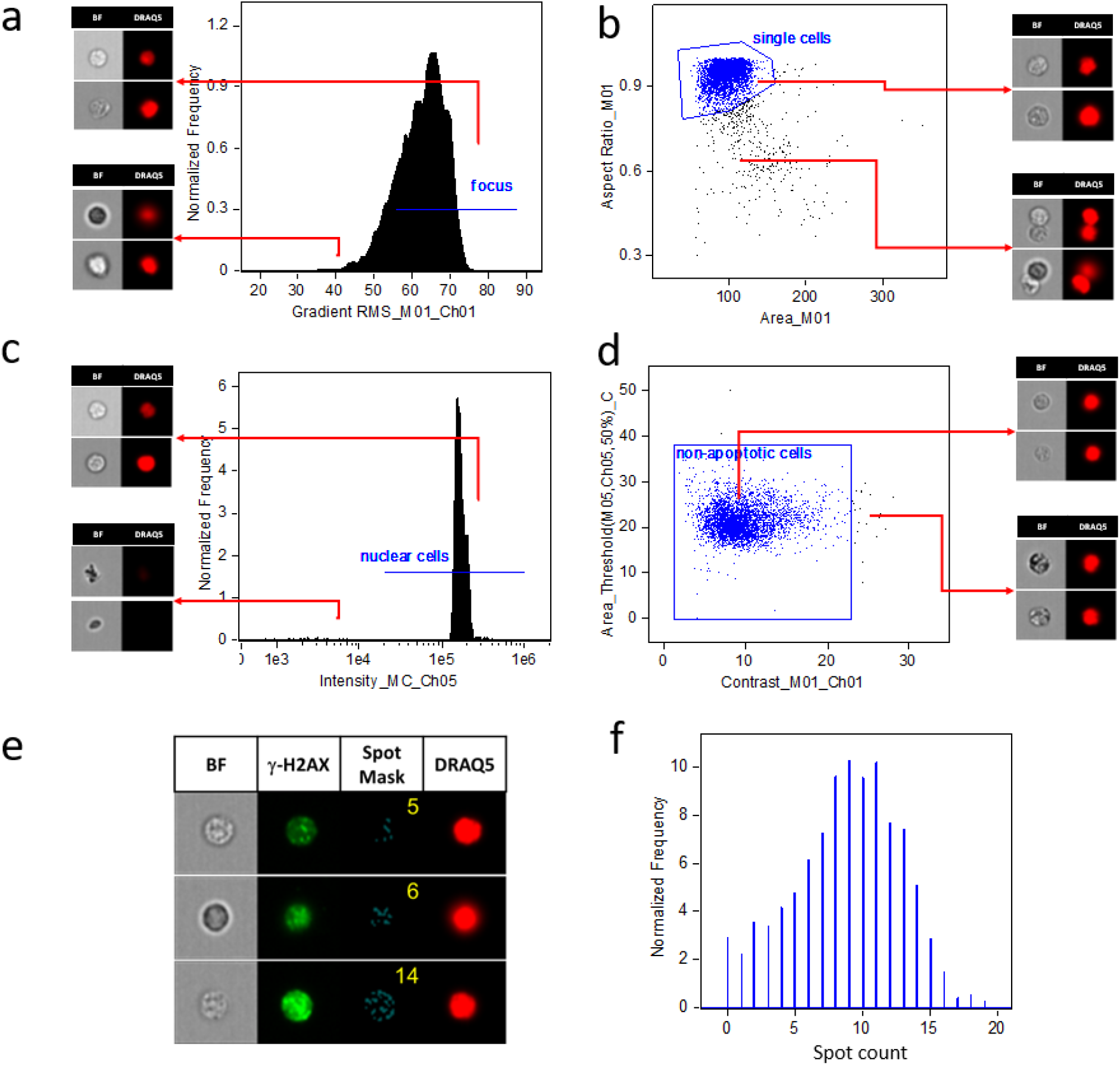
Gating strategy for assessing γ-H2AX levels in the IDEAS® software. (a) According to the gradient similarity feature in the bright field channel, the focused cells were gated; (b) According to area and aspect ratio in the bright field channel, single cells were selected; (c) Nucleated cells are selected based on DRAQ5 positivity; (d) Non-apoptotic cells were gated according to automated image analysis based on nuclear imagery features in combination with bright field morphology. (e) Representative images of γ-H2AX foci analysis in mouse peripheral blood leukocytes. Images display cells on bright field, γ-H2AX staining, γ-H2AX foci mask, DRAQ5 nuclear staining (40X magnification). The spot counts as shown in the spot mask column is automatically enumerated by spot counting wizard. (f) Histogram of spot count performed by wizard in the IDEAS^®^ software to identify the number of γ-H2AX foci per sample. BF=Bright field.

### Statistical analysis

Analyses were performed by GraphPad Prism version 6.00 for Windows (GraphPad Software Inc., La Jolla, CA). The Kruskal-Wallis test was used to compare data among all study groups and the Mann-Whitney U test was then used to compare between irradiated and control groups. The Spearman’s rank correlation coefficient was used to assess accumulated dose dependence of the γ-H2AX level. γ-H2AX’s performance was determined based on Receiver Operating Characteristic (ROC) curves, which allow the characterization of the discrimination between two well-defined populations. The sensitivity, specificity, positive and negative predictive values were evaluated using the optimal threshold value calculated to maximize the Youden’s index. This index is defined as the sum of the sensitivity and specificity (both expressed by a number comprised between 0 and 1) minus 1 to set up the criterion for selecting the optimum cut-off point. All differences were considered statistically significant when p < 0.05.

## Results

### Mouse Dosimetry and Nominal dose

Fig. 3 shows the calculated dose rate, based on the measured source positions, as well as the cumulative dose for each of the two runs. The vertical dips correspond to VADER openings to extract mice.

**Fig. 3:**
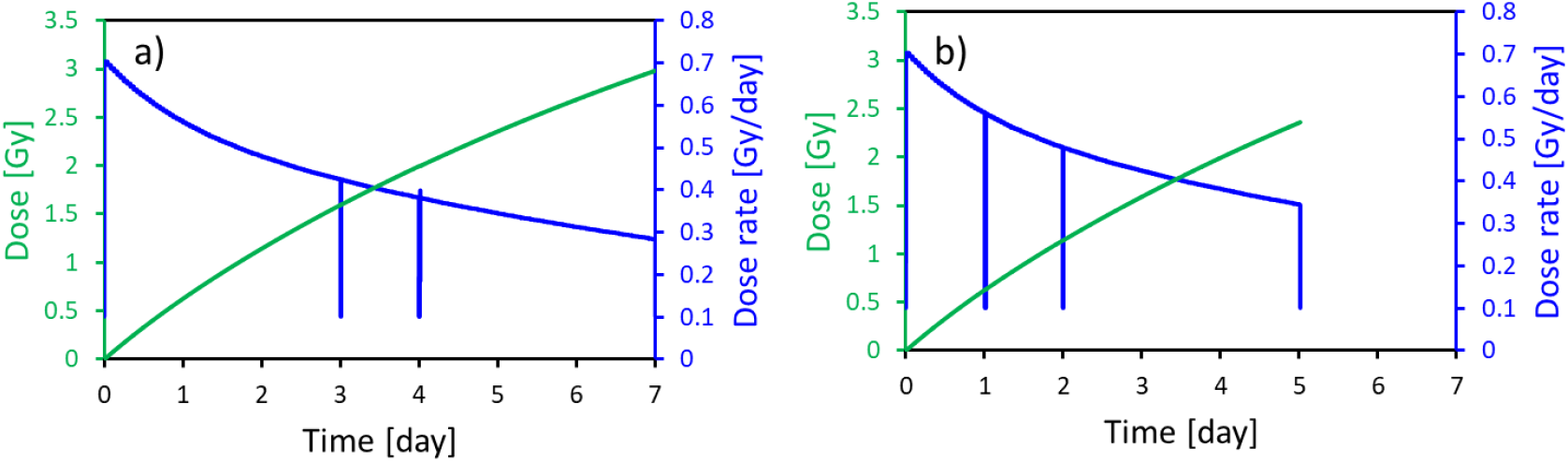
Dose rate (blue), based on the measured source positions, and cumulative doses (green). The dips in the dose rate curve correspond to VADER openings. a) First batch of mice (day 3, day 4 and day7) b) Second batch of mice (day1, day 2 and day5)

### Peripheral blood cell counts kinetics

The total number of peripheral blood leukocytes (CD45+), T cells (CD3+) and B cells (CD19+) were quantified by flow cytometry (Fig. 4). The total blood leukocytes count was not significantly different from the non-irradiated controls until day 7 in the VADER irradiated group (p<0.01) (Fig. 4a), as were the T-cells counts (p<0.05) (Fig. 4b). B cells responded with increased sensitivity to radiation whereby the number of cells had significantly decreased by day 4 and remained low up to 7 days (Fig. 4c).

**Fig. 4.**
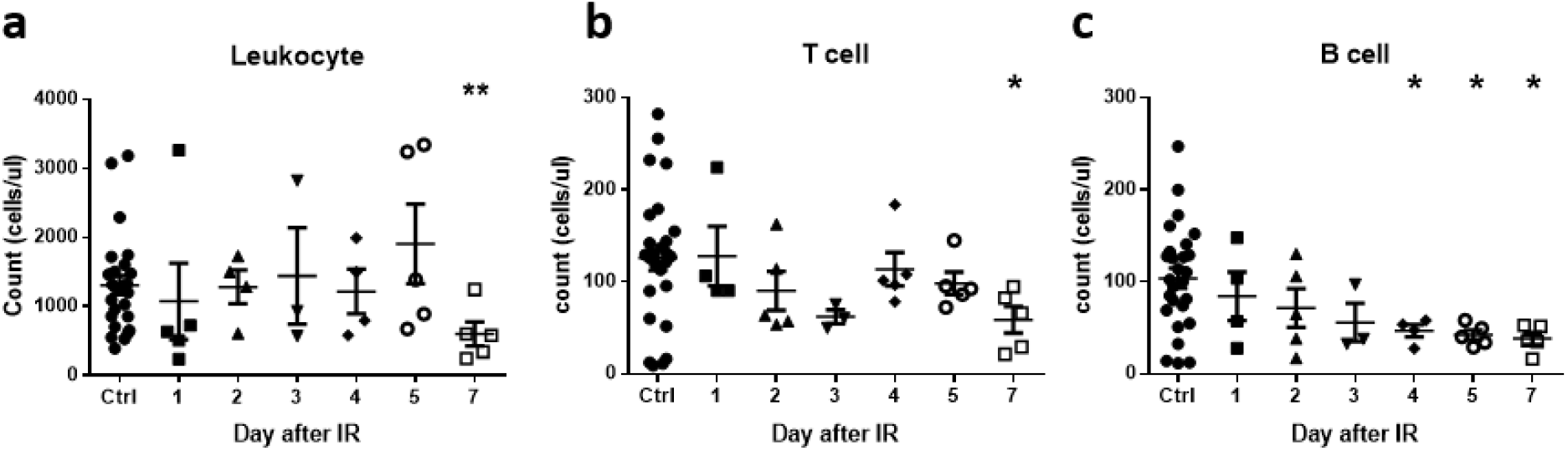
Effects of VADER irradiation on peripheral blood leukocyte, T cell and B cell counts. (a) Leukocyte counts; (b) T cell counts; (c) B cells counts. Error bar is expressed in means ± SEM. The Kruskal-Wallis test was performed to compare data among study groups. The Mann-Whitney U test was then used to compare VADER irradiated group with non-irradiated control group. *p<0.05 and **p<0.01.

### Apoptotic cell counts and P53 expression

The overall rate of apoptotic cells identified by IDEAS^®^ software based on their morphology and nuclear complexity was relatively low (<3%), with no significant difference observed between the VADER irradiation group and the non-irradiated control (data not shown). As p53 is a critical inducer for apoptosis signaling pathway, expression of p53 in the non-apoptotic leukocyte population was also quantified (Fig. 2). To compensate for the random measurement error, the non-irradiated mice p53 MFI were normalized to 1000 MFI units, and the VADER irradiated mice p53 MFI levels were calculated based on the same-day non-irradiated control. The results show that p53 expression peaked on day 2 and then declined, but remained significantly above non-irradiated baseline levels up to day 4 (p<0.05, Fig. 5).

**Fig. 5.**
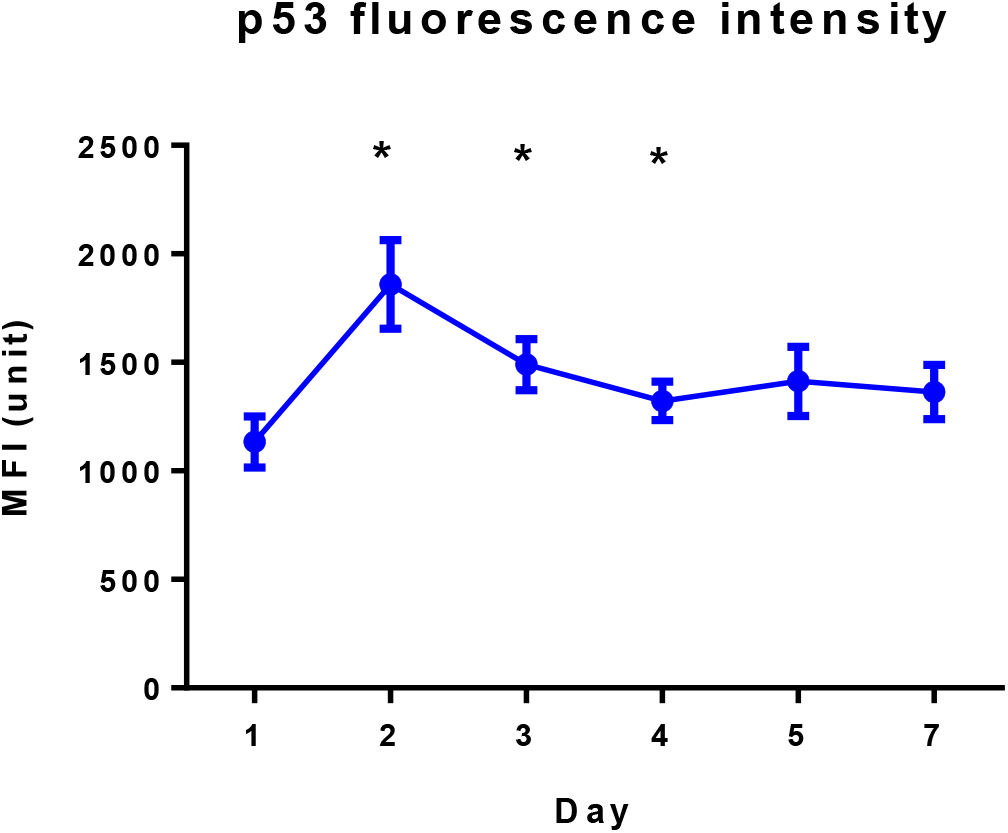
Kinetics of p53 fluorescence levels in peripheral blood leukocytes during VADER irradiation. Error bar is expressed in means ± SEM. The Kruskal-Wallis test was performed to compare data among study groups. The Mann-Whitney U test was used to compare VADER irradiated group with non-irradiated control group. *p<0.05.

### γ-H2AX expression and foci formation

Fig. 6 shows that γ-H2AX fluorescence intensity levels and foci formation displayed a similar trend with increasing accumulated dose during the 7-day study irradiation time. For each time point, γ-H2AX intensity levels of non-irradiated control mice was normalized to the average of 1000 unit of MFI. Foci number was normalized to the average of 5 per cell. The results are presented as the relative MFI and foci number compare with the same day non-irradiated control mice sample. Fig. 6a shows that the γ-H2AX MFI levels is significantly increased by day 3 and peaked at day 5, after which time γ-H2AX levels significantly decreased. Fig. 6b illustrates a similar kinetic curve for γ-H2AX foci formation, where the number of γ-H2AX foci similarly started to increase significantly from day 3 and peaked at day 5.

**Fig. 6.**
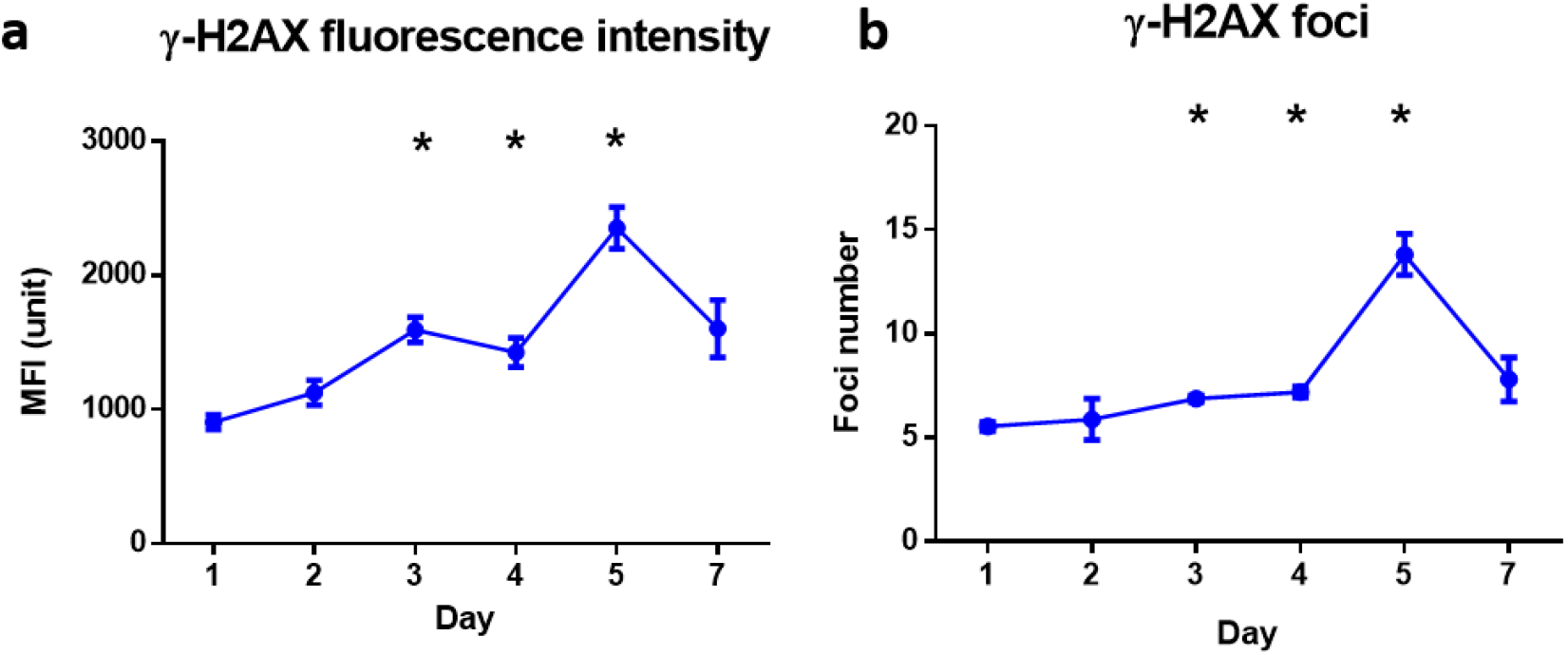
Kinetics of γ-H2AX mean fluorescence intensity (MFI; panel a) and foci number (panel b) in peripheral blood leukocytes during 7 days of VADER exposure. Error bars is expressed in means ± SEM. The Kruskal-Wallis test was performed to compare data among study groups. The Mann-Whitney U test was used to compare VADER irradiated group with non-irradiated control group. *p<0.05.

### γ-H2AX as a biodosimetry marker for the accumulated dose

Since γ-H2AX MFI and foci formation showed a similar trend with accumulated dose, we performed the correlation of accumulated dose with γ-H2AX intensity expression and γ-H2AX foci formation (Fig. 7). The data show that γ-H2AX MFI strongly correlated with accumulated dose (p<0.001, r=0.71) (Fig. 7a), while γ-H2AX foci number moderately correlated with accumulated dose (p<0.001, r= 0.55) (Fig. 7b). Receiver operating characteristic (ROC) analysis was used to evaluate the performance (Lacombe et al. 2018) and estimate the sensitivity and specificity of γ-H2AX MFI to discriminate the “classification threshold” (e.g. 2Gy). The optimal cut off value of γ-H2AX expression level was determined by Youden Index with the highest sensitivity and specificity. The sum value of MFI=1583 units was used as the cut off value to determine whether the accumulated dose is higher than 2 Gy. The area under the curve (AUC) was 0.898 (standard error [SE], 0.061; 95% confidence interval [CI], 0.7761 to 1.019, p < 0.001) (Fig. 7c).

**Fig. 7.**
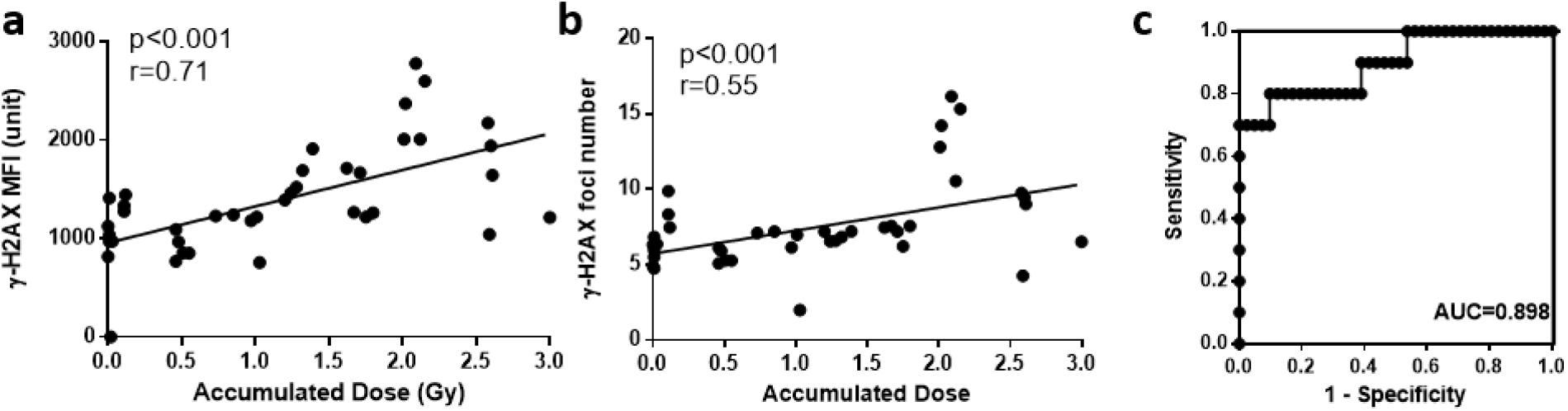
Correlation of gH2AX levels with accumulated dose and performance to discriminate dose. (a) Correlation of the γ-H2AX mean fluorescence intensity with the accumulated dose (p<0.001, r=0.71). (b) Correlation of the γ-H2AX mean fluorescence intensity with the accumulated dose (p<0.001, r=0.55). (c) Receiver operating characteristic (ROC) curve analysis to discriminate radiation dose < 2 Gy from radiation dose ≥ 2 Gy based on γ-H2AX mean fluorescence intensity.

## Discussion

Internal radiation exposure has been considerably understudied largely due to the practicalities and expense associated with ^137^Cs in-vivo studies. The recent development of our ^137^Cs VADER irradiator has allowed us to simulate protracted radionuclide exposures and measure the DNA damage response in mouse peripheral blood leukocytes *in vivo* exposed to progressive, variable low dose-rate gamma rays (Garty et al. 2017). The advantage of the VADER, besides the flexibility in dose and dose rate options is that we were able to collect the biofluids without ^137^Cs radioactive contamination. Besides, by using imaging flow cytometry (IFC), we are able to rapidly measure multiple biomarker endpoints

In the present study, we showed that leukocytes and T cell counts were significantly decreased at day 7, while B cells started to decrease by day 4 (dose = 1.71 Gy) and remained low up to 7 days. These findings are consist with results in previous studies that B cell are more sensitive to irradiation than other peripheral cell population in mouse (Kajioka et al. 2000). Apoptosis/cell death contributes to the depletion of the peripheral leukocytes (Garty et al. 2019b). The use of imaging flow technology has enabled us to acquire the percentage of the apoptotic cells according to their complexity of nucleus and contrast morphology. We found that proportion of apoptotic cells was relatively low (<3%), with no significant difference observed between the VADER irradiation group and the non-irradiated control, whereas p53 levels were significantly above control and increased on day 2, 3 and 4 during VADER irradiation. Functionally, p53 is a transcription factor with control of target genes that influence cell cycle arrest, DNA repair, apoptosis, and senescence (Eriksson and Stigbrand 2010). Depending on the extent of damaged DNA and cell type, p53 can activate elimination routes for damaged cells by apoptosis or senescence (Rodier et al. 2009; Rufini et al. 2013). We speculate that it is possible that in a protracted low dose-rate irradiation scenario, the continuous double-stranded DNA repair activated p53 at early time point, leading to the cell senescence, preventing consequent damage from being propagated to the next cell generation, and finally contribute to the leukocytes depletion at day 7 after irradiation. Recently, Cao. et al demonstrated that the chronic ex-vivo low dose irradiation up to 10 days induced considerably more cellular senescence in human tumor cells lines (Cao et al. 2014).

In contrast to p53 MFI which peaks at day 2, the γ-H2AX MFI and foci formation showed an significant increase with accumulated dose up to day 5, after which levels decreased. ROC analysis showed that γ-H2AX MFI value provide a good discrimination of accumulated dose < 2 Gy and ≥ 2 Gy, suggesting that γ-H2AX is a candidate biomarker for dosimetry in a protracted, environmental exposure scenario. This is consist with recent studies suggested γ-H2AX and 53BP1 could be useful biomarkers for detecting low dose radiation after internal ^131^I injection in thyroid cancer patients (Eberlein et al. 2016; Lassmann et al. 2010).

For development radiation biodosimetry assays, rapid and accurate dose estimations are highly desired. Here, we adopted imaging flow cytometry (IFC) technology for the γ-H2AX and p53 assay. IFC is relatively new technology that has been developed to combine the statistical power of traditional flow cytometry with the sensitivity and specificity of microscopy acquisition speeds of 1000 cells/seconds (Wang et al. 2019). It also allows to analyze cellular morphology and quantitate multi parametric fluorescent intensities (Vorobjev and Barteneva 2016). The application of multispectral imaging flow cytometry provides a novel and robust methodology for the estimation and quantitation of γ-H2AX intensity as well as foci in cells (Lee et al. 2019).

In summary, we have used the novel imaging flow cytometry to develop a rapid γ-H2AX assay for the detection of DNA DSBs in peripheral blood leukocytes after exposure to variable low dose-rate external ^137^Cs irradiator over 7 days. Results present here highlighted the potential of γ-H2AX as a biomarker to provide a good discrimination of accumulated dose < 2 Gy and ≥ 2 Gy in a protracted, environmental exposure scenario. Follow-up studies for biodosimetry measurements using our VADER approach to mimic the low dose radionuclides exposure in accidental or malicious radiological scenario is needed.

## Reference

Cao L et al. (2014) A novel ATM/TP53/p21-mediated checkpoint only activated by chronic γ-irradiation PLoS One 9:e104279

Eberlein U et al. (2016) DNA Damage in Peripheral Blood Lymphocytes of Thyroid Cancer Patients After Radioiodine Therapy Journal of nuclear medicine: official publication, Society of Nuclear Medicine 57:173–179 doi:10.2967/jnumed.115.164814

Eriksson D, Stigbrand T (2010) Radiation-induced cell death mechanisms Tumor Biology 31:363–372

Garty G, Pujol MC, Brenner DJ (2019a) An injectable dosimeter for small animal irradiations arXiv preprint arXiv:190407311

Garty G, Xu Y, Elliston C, Marino SA, Randers-Pehrson G, Brenner DJ (2017) Mice and the A-bomb: irradiation systems for realistic exposure scenarios Radiation research 187:475–485

Garty G et al. (2019b) VADER: a VAriable Dose-rate External 137Cs irradiatoR for internal emitter and low dose rate studies arXiv preprint arXiv:190504169

Goudarzi M et al. (2013) Development of urinary biomarkers for internal exposure by cesium-137 using a metabolomics approach in mice Radiation research 181:54–64

Halm BM, Franke AA, Lai JF, Turner HC, Brenner DJ, Zohrabian VM, DiMauro R (2014) γ-H2AX foci are increased in lymphocytes in vivo in young children 1 h after very low-dose X-irradiation: a pilot study Pediatric radiology 44:1310–1317

Kajioka EH et al. (2000) Acute effects of whole-body proton irradiation on the immune system of the mouse Radiation research 153:587–594

Lacombe J, Sima C, Amundson SA, Zenhausern F (2018) Candidate gene biodosimetry markers of exposure to external ionizing radiation in human blood: A systematic review PloS one 13:e0198851

Lassmann M, Hanscheid H, Gassen D, Biko J, Meineke V, Reiners C, Scherthan H (2010) In vivo formation of gamma-H2AX and 53BP1 DNA repair foci in blood cells after radioiodine therapy of differentiated thyroid cancer Journal of nuclear medicine: official publication, Society of Nuclear Medicine 51:1318–1325 doi:10.2967/jnumed.109.071357

Lee Y, Wang Q, Shuryak I, Brenner DJ, Turner HC (2019) Development of a high-throughput γ-H2AX assay based on imaging flow cytometry bioRxiv:637371

López-Vicente M, Onda Y, Takahashi J, Kato H, Chayama S, Hisadome K (2018) Radiocesium concentrations in soil and leaf after decontamination practices in a forest plantation highly polluted by the Fukushima accident Environmental Pollution 239:448–456

Moysich KB, Menezes RJ, Michalek AM (2002) Chernobyl-related ionising radiation exposure and cancer risk: an epidemiological review The Lancet Oncology 3:269–279

Nakamura AJ et al. (2017) The causal relationship between DNA damage induction in bovine lymphocytes and the Fukushima Nuclear Power Plant Accident Radiation research 187:630–636

Paul S, Ghandhi SA, Weber W, Doyle-Eisele M, Melo D, Guilmette R, Amundson SA (2014) Gene expression response of mice after a single dose of 137CS as an internal emitter Radiation research 182:380–389

Rodier F et al. (2009) Persistent DNA damage signalling triggers senescence-associated inflammatory cytokine secretion Nature cell biology 11:973

Rufini A, Tucci P, Celardo I, Melino G (2013) Senescence and aging: the critical roles of p53 Oncogene 32:5129

Simon SL, Bouville A, Beck HL (2004) The geographic distribution of radionuclide deposition across the continental US from atmospheric nuclear testing Journal of Environmental Radioactivity 74:91–105

Solier S, Pommier Y (2014) The nuclear gamma-H2AX apoptotic ring: implications for cancers and autoimmune diseases Cellular and molecular life sciences: CMLS 71:2289–2297 doi:10.1007/s00018-013-1555-2

Steinhausler F (2005) Chernobyl and Goiania lessons for responding to radiological terrorism Health physics 89:566–574

Turner H et al. (2019) Effect of dose and dose rate on temporal γ-H2AX kinetics in mouse blood and spleen mononuclear cells in vivo following Cesium-137 administration. BMC Molecular and Cell Biology

Turner HC et al. (2010) Adapting the γ-H2AX assay for automated processing in human lymphocytes. 1. Technological aspects Radiation research 175:282–290

Turner HC et al. (2015) gamma-H2AX Kinetic Profile in Mouse Lymphocytes Exposed to the Internal Emitters Cesium-137 and Strontium-90 PloS one 10:e0143815 doi:10.1371/journal.pone.0143815

Vorobjev IA, Barteneva NS (2016) Quantitative Functional Morphology by Imaging Flow Cytometry Methods in molecular biology 1389:3–11 doi:10.1007/978-1-4939-3302-0_1

Wang Q et al. (2019) Automated Triage Radiation Biodosimetry: Integrating Imaging Flow Cytometry with High-Throughput Robotics to Perform the Cytokinesis-Block Micronucleus Assay Radiation research

Wang Q et al. (2016) Reduced levels of cytosolic DNA sensor AIM2 are associated with impaired cytokine responses in healthy elderly Experimental gerontology 78:39–46

Yasunari TJ, Stohl A, Hayano RS, Burkhart JF, Eckhardt S, Yasunari T (2011) Cesium-137 deposition and contamination of Japanese soils due to the Fukushima nuclear accident Proceedings of the National Academy of Sciences 108:19530–19534

